# Gene editing of *CD3 epsilon* gene to redirect regulatory T cells for adoptive T cell transfer

**DOI:** 10.1101/2024.03.18.584896

**Authors:** Weijie Du, Fatih Noyan, Oliver McCallion, Vanessa Drosdek, Jonas Kath, Viktor Glaser, Carla Fuster-Garcia, Mingxing Yang, Maik Stein, Olaf Weber, Julia K. Polansky, Toni Cathomen, Elmar Jaeckel, Joanna Hester, Fadi Issa, Hans-Dieter Volk, Michael Schmueck-Henneresse, Petra Reinke, Dimitrios L. Wagner

## Abstract

Adoptive transfer of regulatory T cells (Tregs) is a promising strategy to combat immunopathologies in transplantation and autoimmune diseases. Antigen-specific Tregs are more effective in modulating undesired immune reactions, but their low frequency in peripheral blood poses challenges for manufacturing and their clinical application. Chimeric antigen receptors (CARs) have been used to redirect the specificity of Tregs, employing retroviral vectors. However, retroviral gene transfer is costly, time consuming, and raises safety issues. Here, we explored non-viral gene editing to redirect Tregs with CARs, using HLA-A2-specific constructs for proof-of-concept studies in transplantation models. We introduce a virus-free CRISPR-Cas12a approach to integrate an antigen-binding domain into the *CD3 epsilon* (*CD3ε*) gene, generating Tregs expressing a T cell receptor fusion construct (TruC). These *CD3ε*-TruC Tregs exhibit potent antigen-dependent activation while maintaining responsiveness to TCR/CD3 stimulation. This enables preferential enrichment of TruC-redirected Tregs via repetitive CD3/CD28-stimulation in a GMP-compatible expansion system. Non-viral gene edited *CD3ε*-TruC Tregs retained their phenotypic, epigenetic, and functional identity. In a humanized mouse model, HLA-A2-specific *CD3ε*-TruC Tregs demonstrate superior protection of allogeneic HLA-A2^+^ skin grafts from rejection compared to polyclonal Tregs. This approach provides a pathway for developing clinical-grade *CD3ε*-TruC-based Treg cell products for transplantation immunotherapy and other immunopathologies.

## II. Introduction

Regulatory T cells (Tregs) are immunosuppressive lymphocytes that prevent autoimmune diseases, modulate and end overshooting immune responses, promote tissue regeneration and regulate bacterial homeostasis at mucosal surfaces. In circulation, 2-5% of the peripheral blood’s CD4^+^ T cells represent Tregs characterized by a canonical phenotype, such as the high expression of the master transcription factor FOXP3 and the IL-2 receptor alpha chain, CD25, as well as low expression of the IL-7 receptor alpha chain, CD127. Clinical data from patients with solid organ or hematopoietic stem cell transplantation associate higher frequency of activated Tregs after transplantation with better graft survival and lower incidence of graft-versus-host-disease, respectively^1–4^. Congenital Tregs dysfunction is associated with severe autoimmune disease. Consequently, adoptive Tregs transfer has emerged as a modality for tailored immunosuppression with many potential applications.

Adoptive transfer of *ex vivo* expanded (1^st^ generation) polyclonal Tregs has been clinically tested as a potential treatment to minimize immunosuppression in solid organ transplants^5^, resolve chronic graft versus host disease^6^, ameliorate autoimmune diseases, such as type 1 diabetes mellitus^7^ or inflammatory bowel diseases^8^. Transfer of antigen-specific Tregs has demonstrated higher potency in most preclinical models of above-mentioned diseases^9^. Challenges in manufacturing arise from the low abundancy of antigen-specific Tregs in the peripheral blood, which has slowed adoption of antigen-specific Tregs in clinical practice^10^.

Chimeric antigen receptors (CARs) are tools to guide Tregs toward target antigens, such as auto-, allo- or tissue-specific antigens. CARs are synthetic fusion genes, which typically combine the antigen-binding domain of a monoclonal antibody (commonly single-chain variable fragment [scFv]) with intracellular signaling domains that trigger T cell activation, usually via CD3 zeta. Conventional 2^nd^ generation CARs incorporate additional co-stimulatory signaling domains to enhance T cell proliferation and survival. In Tregs, CD28 has been favored as a costimulatory domain in 2^nd^ generation CARs^11^.

The first CAR-Treg candidate that entered clinical trial uses an HLA-A2-specific CAR construct^12^ and is tested in allogeneic kidney transplantation^13^ (NCT04817774). Scarcity of donor organs, difficulty of HLA-matching and potent immunosuppressive drug cocktails established minimally HLA-matched allogeneic solid organ transplantation as the gold-standard, in particular in living-donor recipients. Due to the high frequency of the HLA-A2 allele in the Caucasian population, approximately a quarter of HLA-A2-negative renal transplant recipients were offered an HLA-A2 positive renal transplant in Europe and the US^13^. HLA-A2 represents a promising candidate for CAR targeting in transplant medicine^14,15^. Results on safety and efficacy of HLA-A2-directed Tregs in clinical trials are pivotal to inform future developments for autoimmune diseases.

Manufacturing of CAR-redirected Tregs is dominated by retroviral gene transfer (e.g. replication-deficient lentivirus)^14–16,12^. Production of viruses for gene transfer is costly and time consuming at clinical stages. Non-viral approaches with transposase technology, such as *PiggyBac* or *Sleeping Beauty*, represent more affordable alternatives for gene transfer. To our knowledge, they have not been adopted in the Treg space. Furthermore, the two recent cases of CAR-derived lymphoma in a trial using hyperactive PiggyBac modified T cells^17,18^ attest a potential risk of mutagenesis when using transposase technology which may be attributed to random insertion of multiple transgene copies, genetic dysregulation by exogenous promoters or genetic scarring by transgene hopping of transposable elements^19^.

CRISPR-Cas gene editing may represent a non-viral alternative for site-specific gene transfer in Tregs^20^, which reduces the risk for insertional mutagenesis. By utilizing genomic promoters, it may further alleviate the need for exogenous promoters for efficient transgene expression. Virus-free knock-in through electroporation of dsDNA and CRISPR-Cas9 ribonucleoprotein (RNP) complexes was shown to allow efficient reprogramming of T cells and Tregs ^20–23^. Studies using this approach usually integrate the transgenic antigen receptor into the TCR alpha constant gene (*TRAC*), because this leads to TCR replacement with low risk of alloreactivity and physiological regulation of the transgene, which has been associated with improved CAR-redirected effector T cell fitness in leukemia models^24,25^. However, optimal *ex vivo* expansion of Treg requires distinct stimuli and metabolic conditions from conventional CAR-T cells used for cancer treatment limiting a 1:1 knowledge transfer to optimal CAR-Treg design.

In this study, we evaluated virus-free knock-in of CARs to redirect Tregs and used HLA-A2 specific CAR for functional proof-of-concept studies. As TCR/CD3-negative *TRAC*-replaced CAR Tregs failed to expand efficiently, we devised a novel promoter less gene editing strategy to produce redirected Tregs in a virus-free and GMP-compatible process. CRISPR-Cas12a mediated insertion of a fully human HLA-A2-specific scFv cDNA into the *CD3 epsilon* (*CD3ε*) gene facilitated the generation of T cell receptor fusion construct (TruC)-expressing human Tregs. Functional characterization *in vitro* and *in vivo* highlights the potent immunosuppressive properties of *CD3ε*-TruC Tregs.

## III. Results

### Identification of a CD3 epsilon gene editing strategy to create TruC+ T cells after integration of scFv cDNA

To generate an HLA-A2-specific CAR construct for redirecting Tregs, we replaced the antigen-binding domain of 2^nd^ generation CAR^23^ with an HLA-A2-specific scFv which was originally derived from an allo-sensitized 57-year-old female patient^26^ (**Suppl. Fig. 1 A, B**). Following PBMC isolation from peripheral blood of HLA-A2-negative healthy donors, CD4^+^CD25^high^CD127^low^CCR7^+^ Tregs were sorted with more than 90% of purity (**Suppl. Fig. 2**). Then, Tregs were activated using anti-CD3/28 stimulation beads and expanded in the presence of IL-2 and the mTOR inhibitor rapamycin^27^. After 7-9 days expansion, the proliferating Tregs were harvested and electroporated with linear double stranded (ds)DNA for CAR integration via homology-dependent DNA-repair (HDR-template) and a pre-complexed RNP containing *TRAC*-specific guide RNA (gRNA)^23^, recombinant *Streptococcus pyogenes* (S.p.) Cas9 protein and poly-L-glutamic acid (PGA)^28^. The resulting *TRAC*-CAR^+^ Tregs were predominantly CD3-negative (**Suppl. Fig. 1C**) and retained canonical markers of Tregs identity e.g. high expression of FOXP3 and CD25 (data not shown). However, *TRAC-*CAR Tregs could not be efficiently expanded after gene editing either via HLA-A2 overexpressing K562 cells or dimeric human HLA-A2:Ig fusion protein stimulation (**Suppl. Fig. 1 D**). Re-stimulation on plate-bound recombinant HLA-A2:Ig-fusion protein every 2-3 days with and without anti-CD28 mAb yielded expansion folds of 1.60 and 2.57 respectively, after gene editing. Addition of irradiated HLA-A2 overexpressing K562 cells improved expansion from 0.5-fold (no stimulation) up to 5-fold. In contrast, polyclonally expanded Tregs could expand up to 500-fold within a 19-day expansion period (**Suppl. Fig. 1 D**).

The suboptimal expansion of *TRAC*-CAR Tregs is prohibitive to create a stable protocol at clinical scale. Therefore, we devised a gene editing strategy to redirect Tregs with the chosen HLA-A2-specific scFv but compatible with our GMP expansion process using repetitive anti-CD3/28 bead stimulation^5^. To this end, we hypothesized that targeted insertion of the scFv cDNA into the endogenous *CD3 epsilon* (*CD3ε*) gene should generate HLA-A2-specific T cells which express a T cell receptor fusion construct (TruC) consisting of transgenic antigen-binding domain and the endogenous CD3ε protein. Due to the ability of TruC to form complex with the endogenous TCR/CD3 complex^29^, Tregs with the TruC knock-in strategy should retain TCR/CD3 complex on their cell surface, which can be engaged via anti-CD3/28 stimulation beads as well as the HLA-A2 antigen.

Homology-directed repair with programmable nucleases depends on efficient installation of DNA double strand breaks and the subsequent occurrence of insertion and deletions. Thus, we first screened 5 potential crRNA candidates (**Suppl. Table 1**) for the CRISPR-Cas12a nuclease derived from *Acidaminococcus species* (AsCas12a) for their ability to disrupt *CD3ε* in exon 3 or exon 6 in conventional T cells (Tconv) (**Fig. 1B**). Synthetic crRNAs and recombinant AsCas12a Ultra^30^ were co-electroporated in polyclonally activated Tconv, and CD3ε expression was measured by flow cytometry as a readout. Four days after nucleofection, Tconv treated with AsCas12a crRNA #1 and crRNA #5 displayed highest KO efficiencies compared to other crRNAs in their respective target sites (**Fig. 1 A, C, D**).

**Figure 1.**
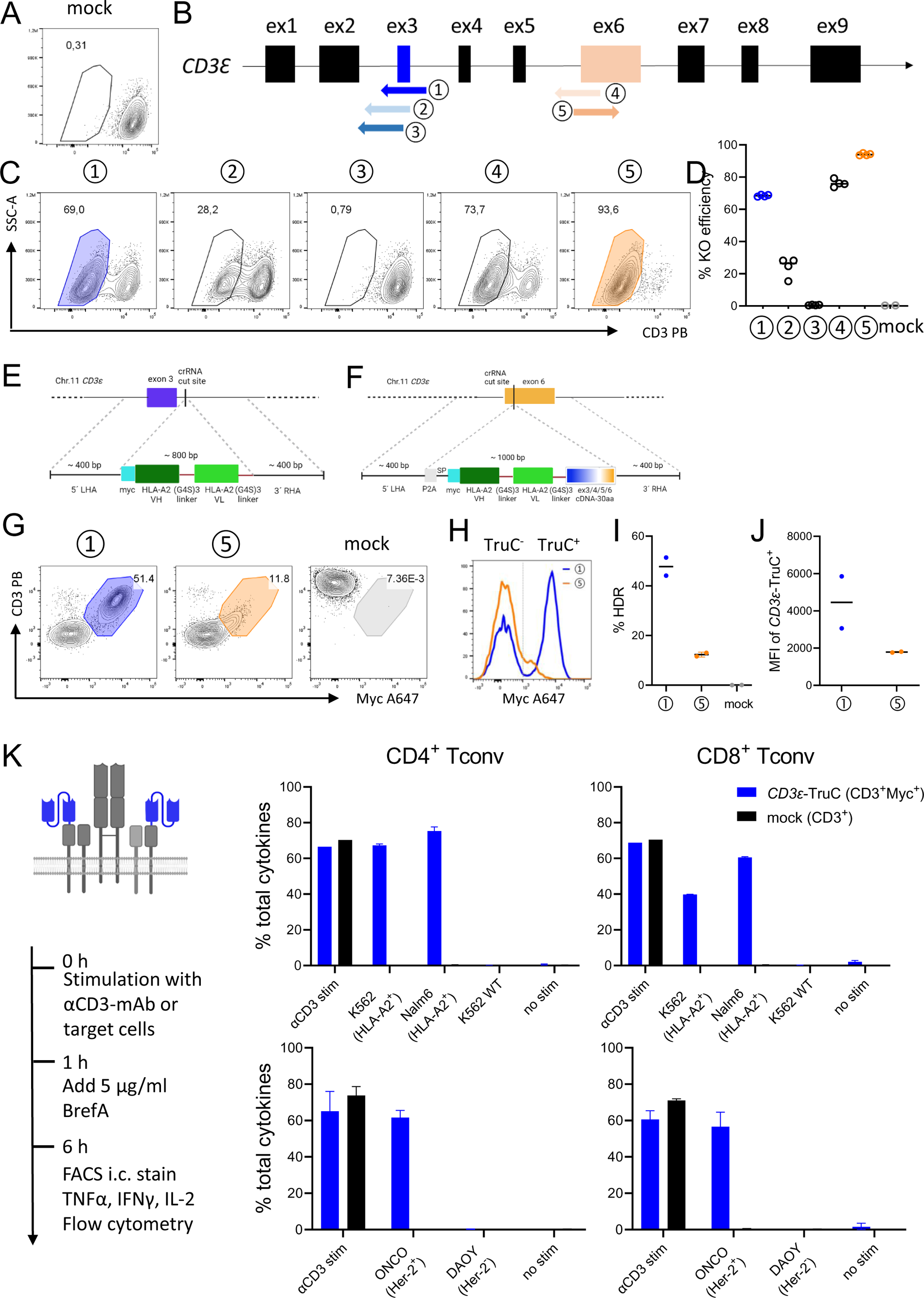
Identification of a CD3 *epsilon* gene editing strategy to create TruC+ T cells after integration of scFv cDNA. A) Representative dot plot of mock electroporated T cells. B) Five crRNAs targeting on exon 3 and exon 6 of CD3 *epsilon* locus were selected. C) Representative dot plots of T cells upon RNP-mediated gene knock-out (KO). D) KO efficiency summary using the five crRNAs. N=4 for KO condition, N=2 for mock condition. HLA-A2 scFv incorporated *CD3ε*-TruC construct design and integration into E) crRNA #1 cut site within intron 3 and into F) crRNA #5 cut site within exon 6. G) Representative dot plots of HLA-A2 *CD3ε*-TruC detection by using crRNA #1 (blue) and crRNA #5 (orange) with mock T cells as staining control (gray). H) Histogram overlay of Myc^+^ *CD3ε*-TruC cells using either crRNA #1 or crRNA #5 for KO. I) HDR efficacy comparison for HDR-templates integrated in crRNA #1 and crRNA #5 cut site, respectively. J) MFI comparison of *CD3ε*-TruC+ cells by integrating TruC construct into *CD3Ε* exon 3 and exon 6, respectively. K) Cytokine profiles of HLA-A2 *CD3ε*-TruC CD4^+^ Tconv (upper left) and CD8^+^ Tconv (upper right) as well as HER2 *CD3ε*-TruC CD4^+^ Tconv (lower left) and CD8^+^ Tconv (lower right) upon anti-CD3 antibody and respective antigen-expressing tumor cell stimulation.

For the two most efficient crRNAs (#1 and #5, targeting exon 3 and exon 6, respectively), we designed corresponding HDR-templates with 400 bp homology arms flanking the HLA-A2-scFv cDNA and the G4S linker, to facilitate the attachment of the antigen-binding domain onto the CD3ε protein (**Fig. 1 E+ F**). For detection purposes, we added a myc-tag in front of scFv cDNA. After transfection of RNP and the respective dsDNA HDR-template into Tconv via electroporation, we could detect large fractions of myc^+^ TruC cells (**Fig. 1 G**). TruC knock-in rates and expression level were significantly higher when targeting exon 3 (**Fig. 1 H, I, J**). To further examine cutting efficiency by Cas12a RNP with crRNA #1, genomic DNA was extracted from two-week expanded *CD3ε*-KO or mock-electroporated Tconv, a T7E1 assay and Sanger sequencing confirmed more than 80% indel frequency in the *CD3ε-*KO Tconv in all three biological replicates, suggesting effective cutting of the CRISPR-AsCas12a RNP with crRNA #1 (**Suppl. Fig. 3**). Thus, we selected crRNA #1 and the respective HDR-template targeting *CD3ε* exon 3 for all future experiments.

To confirm that this *CD3ε*-TruC gene editing strategy enables T cells to signal via *CD3ε*-TruC antigen-binding domain and the TCR/CD3 complex, we generated HLA-A2-specific and HER2-specific CD3ε-TruC Tconv and evaluated their abilities to produce cytokines upon co-cultures with antigen-expressing tumor cells or anti-CD3 stimulation (**Fig. 1 K**). Both HLA-A2- and HER2-specific *CD3ε*-TruC Tconv showed strictly antigen-dependent cytokine production after co-cultures with HLA-A2^+/-^ or HER2^+/-^ tumor cells. These two *CD3ε*-TruC Tconv showed similar levels of cytokine production as CD3^+^ mock Tconv upon pan-T cell stimulation with plate-bound anti-human CD3 antibody. Further, the HLA-A2-specific *CD3ε*-TruC Tconv secreted similar amounts of cytokines as HER2-redirected *CD3ε*-TruC Tconv (**Fig. 1 K**). Collectively, *CD3ε*-TruC strategy allowed dual stimulation through both antigen-scFv interaction and the TCR/CD3 complex in Tconv.

### Off-target analysis with CAST-Seq confirms high specificity of the crRNA-AsCas12a gene editing

The risk of off-target effects and other forms of genotoxicity is a concern when translating CRISPR-Cas to clinical protocols^31^. To confirm high specificity of our preferred CRISPR-AsCas12a gene editing strategy, we examined the off-target profile of the selected AsCas12a-crRNA #1 using CAST-Seq technology^32^ which detects potential off-target sites by identification of translocations between the on-target and the putative off-target sites. CAST-Seq revealed no off-target-mediated translocations (OMTs), suggesting high specificity (**Fig. 2 A, Suppl. Table 4**). At crRNA #1 cleavage site, CAST-Seq revealed the expected pattern of larger insertions and deletions specifically in RNP-treated *CD3ε*-KO Tconv (**Fig. 2 B**). Consequently, our preclinical analysis supports the further development of AsCas12a-crRNA #1 combination for clinical cell products of *CD3ε*-TruC T cells.

**Figure 2.**
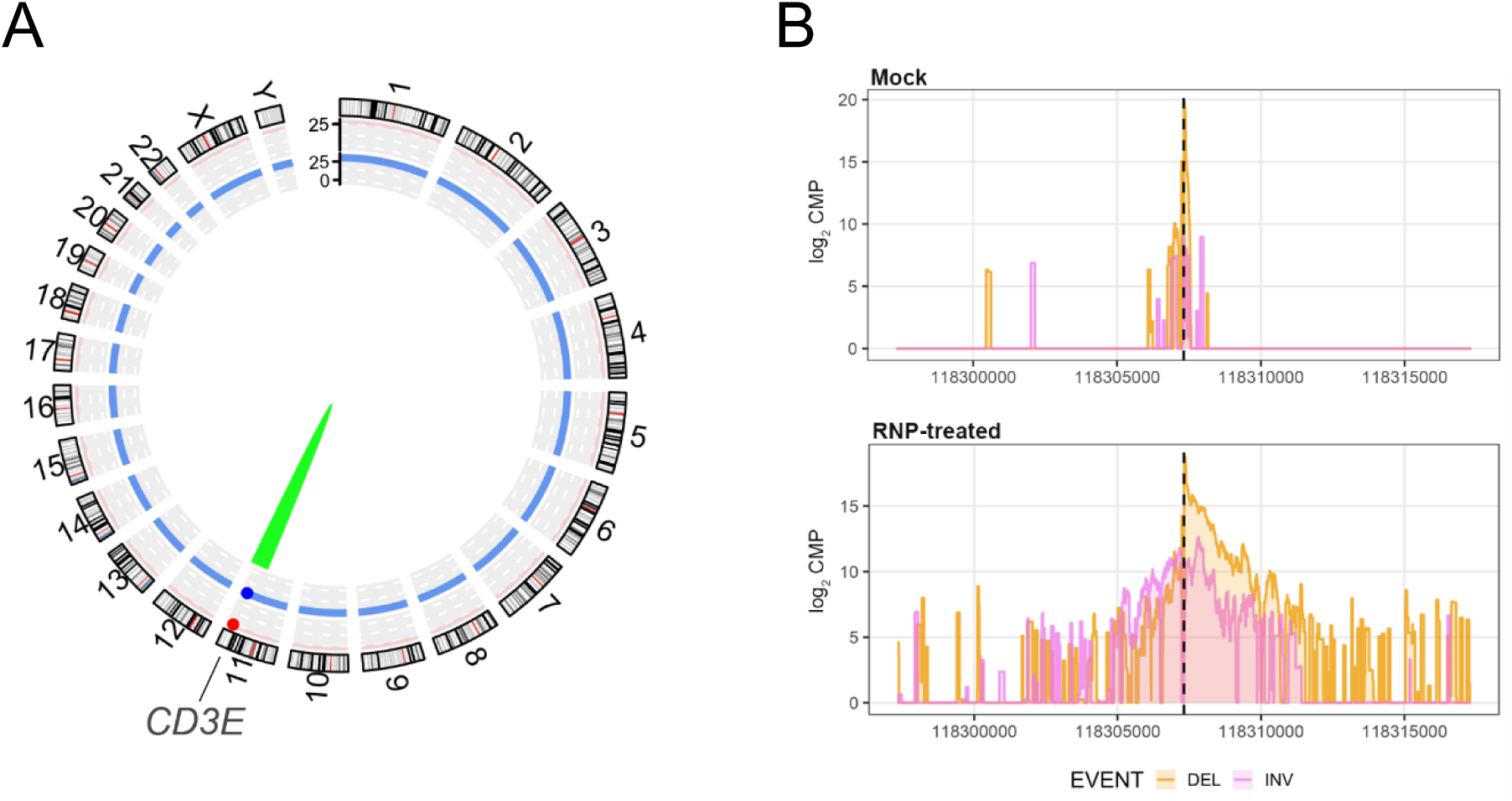
Off-target analysis with CAST-Seq confirms high specificity of the crRNA-Cas12a targeting exon 3 of *CD3Ε*. A) Structural variations. Genomic aberrations at the on-target site are displayed in green, and chromosomal translocations with off-target sites in red (not detected). B) Chromosomal rearrangements at the on-target site. Shown are CAST-Seq coverage plots of reads aligned to a ±10 kb region flanking the *CD3ε* target site. Deletions (DEL) are shown in orange, inversions (INV) in purple. The x-axis represents the chromosomal coordinates, the y-axis the log2 read count per million (CPM), and the dotted line the cleavage site. Sequencing direction is from left to right.

### Continuous CD3/CD28-stimulation leads to preferential expansion of CD3ε-TruC Tregs over TCR-negative Tregs

Using our non-viral gene editing protocol for Tregs, we generated HLA-A2-specific *CD3ε*-TruC, *TRAC*-CAR Tregs and wildtype (WT), unedited Tregs. Following generation, the cells underwent further expansion for two weeks via anti-CD3/28 coated bead stimulation every 2 to 3 days (**Fig. 3 A**). Gene-edited Tregs exhibited a temporary delay in cell growth within the initial 4 days post-electroporation, potentially attributable to the toxicity of the electroporation procedure itself and the use of dsDNA templates. Subsequently, *CD3ε*-TruC Tregs demonstrated robust expansion (**Fig. 3 B, C**). Surprisingly, despite CRISPR-mediated loss of the TCR/CD3-complex, *TRAC*-CAR Tregs also expanded during anti-CD3/28 bead stimulation, albeit at a lower rate than *CD3ε-TruC* Tregs (**Fig. 3 B, C**). Although gene editing of *CD3ε* was less efficient in Tregs than in Tconv (**Suppl. Fig. 4**), *CD3ε*-TruC knock-in rates were comparable to *TRAC*-CAR knock-in rates in the same donor (**Fig. 3 D**). During expansion with anti-CD3/28 bead stimulation, we observed a preferential expansion of CD3^+^ cells, including *CD3ε*-TruC Tregs, over CD3 negative Tregs (**Fig. 3 E**). In contrast to the *CD3ε*-TruC, *TRAC*-CAR^+^ Tregs slightly decreased in frequency along expansion. Similar to the relative transgene^+^ frequency, absolute fold expansion of gene-edited HLA-A2-redirected Tregs was higher in *CD3ε*-TruC Tregs compared to their *TRAC*-CAR counterparts (**Fig. 3 F, G**).

**Figure 3.**
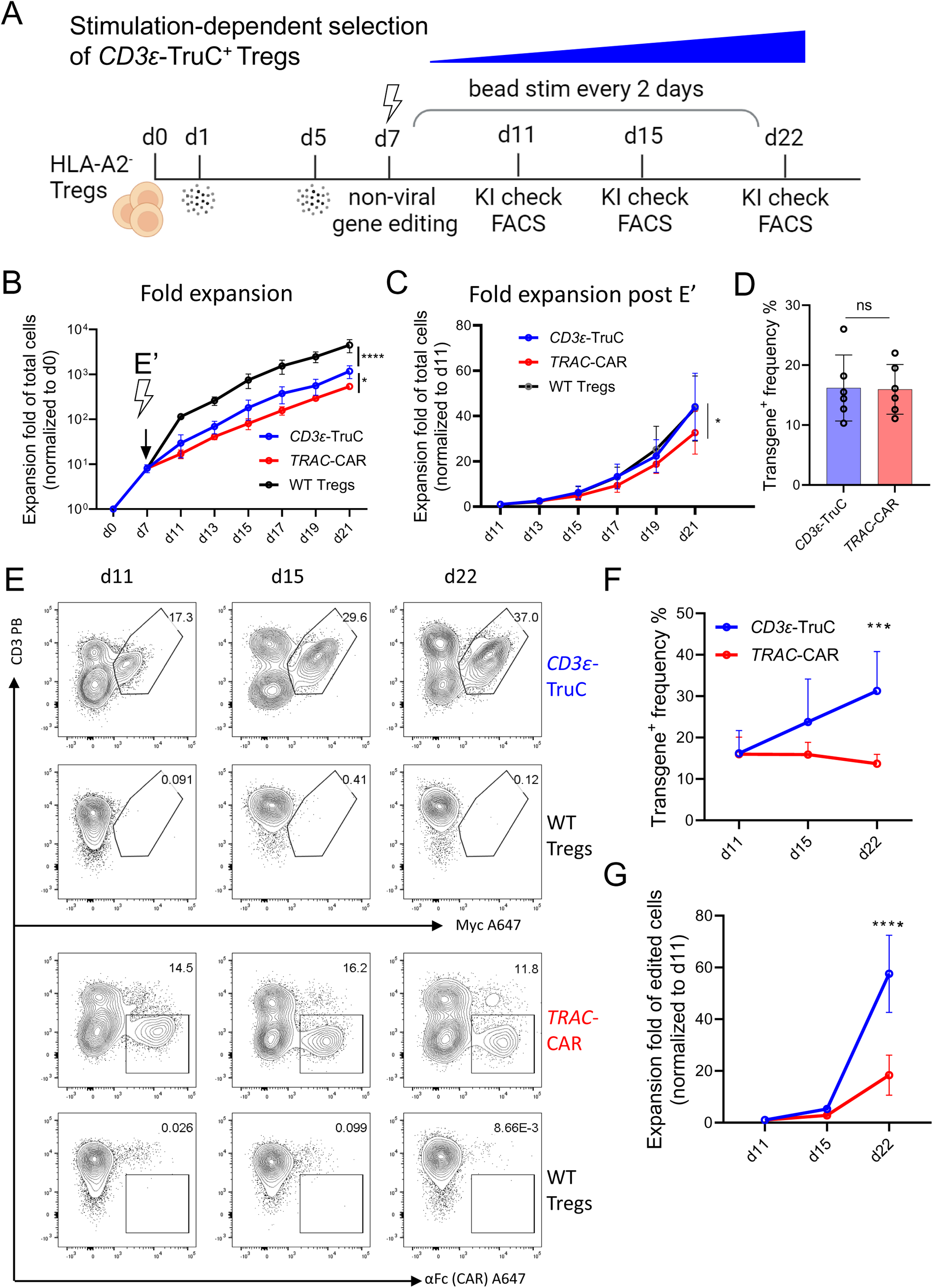
Continuous CD3/CD28-stimulation leads to preferential expansion of *CD3ε*-TruC Tregs over TCR-negative Tregs. A) Schematic pipeline of *CD3ε*-TruC and *TRAC*-CAR Tregs generation and expansion. B) Expansion fold change of total viable cells of *CD3ε*-TruC, *TRAC*-CAR and WT Tregs, normalized to d0. N=4. C) Expansion fold change of total viable cells of *CD3ε*-TruC, *TRAC*-CAR and WT Tregs post electroporation, normalized to d11. N=4. D) Transgene frequencies of *CD3ε*-TruC Tregs and *TRAC*-CAR Tregs at day 11. N=6. E) Representative dot plots of *CD3ε*-TruC (upper) and *TRAC*-CAR (lower) Tregs along expansion. F) Transgene frequencies of *CD3ε*-TruC and *TRAC*-CAR Tregs along expansion. N=6. G) Expansion fold change of gene-edited *CD3ε*-TruC and *TRAC*-CAR positive Tregs, normalized to d11. N=4. Statistical analysis in B, C, F and G was performed using an ordinary two-way ANOVA of matched data. Multiple comparisons were performed by comparing each cell mean with the other cell mean in that row with Turkey (B, C) or Šidák (F, G) correction. Asterisks represent adjusted *p*-values calculated in the respective statistical tests (*: *p* < 0.05; **: *p* < 0.01; ***: *p* < 0.001; ****: *p* < 0.0001). Statistical analysis in D was performed using a t-tests of paired data. Asterisks represent two-tailed *p*-value (ns: *p* ≥ 0.05).

After a two-week expansion post-electroporation, *CD3ε*-TruC, *TRAC*-CAR Tregs retained the canonical Treg phenotype, with more than 98% of total live Tregs express high levels of CD25 and FOXP3, similar to unedited WT Tregs. (**Fig. 4 A, B**). Further, untouched and gene edited Tregs did not secrete Th1 cytokines, e.g. IFNγ and TNFa, after polyclonal restimulation (**Fig. 4 C, D**). Analysis of the Treg-specific demethylated region (TSDR) in the *FOXP3* gene locus was performed to confirm that Tregs retained epigenetic identity with or without gene editing (**Fig. 4 E**). In contrast, unedited conventional T cells of the same donors from the same region were highly methylated (**Fig. 4 E**).

**Figure 4.**
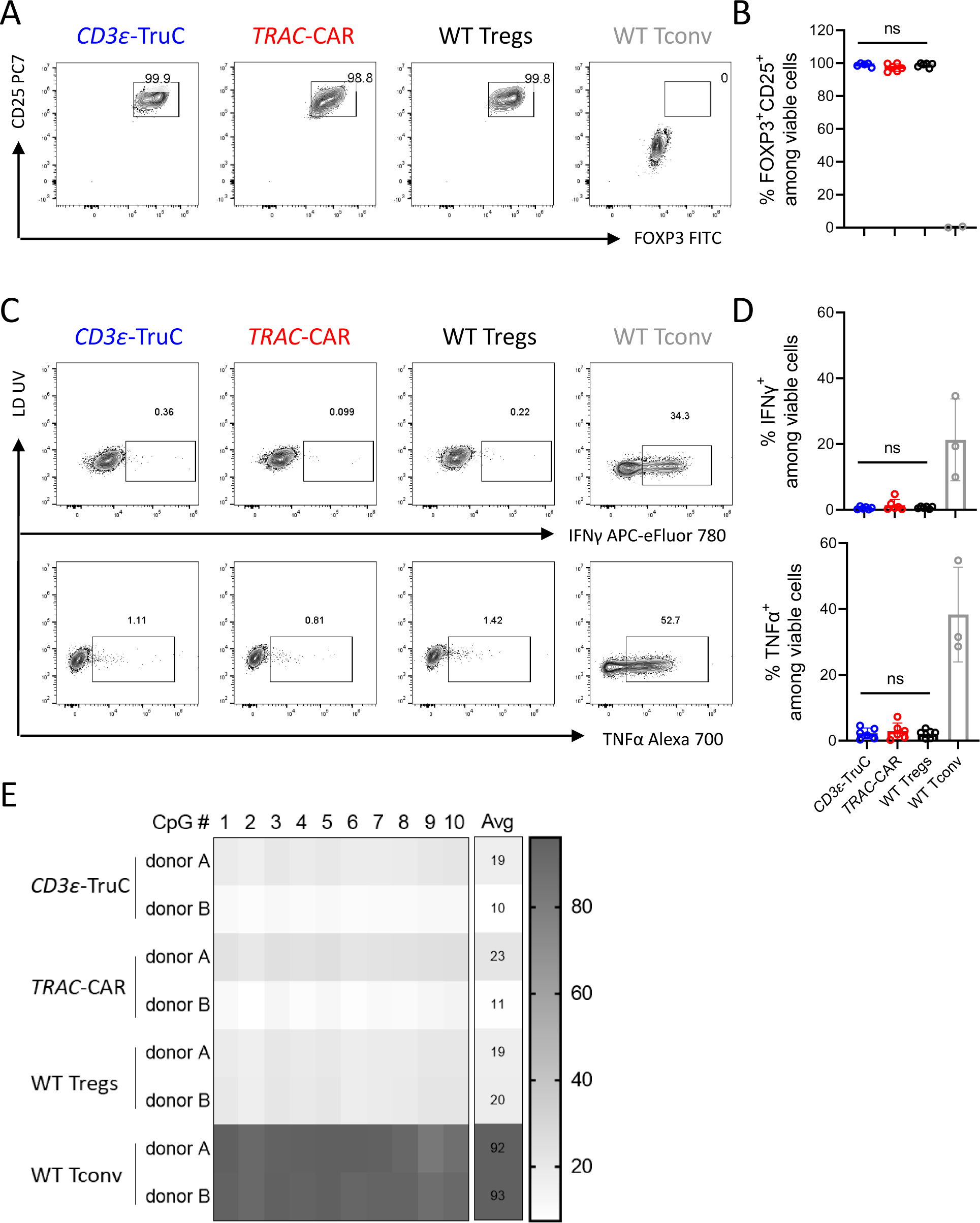
Gene-edited and *ex vivo* expanded Tregs retain their canonical phenotypic and epigenetic identities. A) Representative dot plots of FOXP3 against CD25 among *CD3ε*-TruC, *TRAC*-CAR, WT Tregs and WT Tconv. B) Percentages of FOXP3 and CD25 double positive cells of *CD3ε*-TruC Tregs compared to *TRAC*-CAR and WT Tregs as well as Tconv. N=4. C) Representative dot plots and D) expression level of cytokines IFNγ and TNFα of *CD3ε*-TruC, *TRAC*-CAR and WT Tregs as well as Tconv upon PMA and Ionomycin stimulation. E) Methylation status of 10 individual CpGs at TSDR (Treg specific demethylation region). Values shown in the Avg column indicate the averaged methylation value of 10 CpGs of each sample at TSDR. N=2. Statistical analysis in B and D was performed using a one-way ANOVA of matched data with Geisser-Greenhouse correction. Multiple comparisons were performed by comparing each cell mean with the other cell mean in that row with Turkey correction. Asterisks represent adjusted *p*-value (ns: *p* ≥ 0.05).

### CD3ε-TruC Tregs but not TRAC-CAR Tregs mirror clinically validated polyclonal Tregs

Next, we compared the activation profiles of *CD3ε*-TruC and *TRAC*-CAR Tregs by measuring activation markers in response to different concentrations of plate-bound HLA-A2:Ig fusion protein (**Suppl. Fig. 5 A**). Both *CD3ε-TruC* and *TRAC*-CAR Tregs displayed dose-dependent upregulation of ICOS, CD25, FOXP3, CD137, CD71, LAP and to a limited extend of CTLA4 and Helios (**Suppl. Fig 5 B**). *CD3ε*-TruC displayed a stronger relative (compared to their respective baseline expression without stimulation) upregulation of ICOS, CD25, CD137 compared to *TRAC*-CAR Tregs, especially when stimulated at a higher concentration of 5 µg/ml or 10 µg/ml of HLA-A2:Ig protein (**Suppl. Fig. 5 B**). Of note, *TRAC*-CAR Tregs displayed significantly higher baseline expression of ICOS, CD25, CTLA-4, CD71 and CD137, in comparison to both *CD3ε-* TruC and WT Tregs and may indicate antigen-independent signaling of the CAR (**Suppl. Fig. 5 C**). In contrast, *CD3ε-TruC* and WT Tregs share similar activation profile at baseline without stimulation (**Suppl. Fig. 5 C**).

The non-viral gene editing protocol enabled us to manufacture *CD3ε*-TruC Tregs more efficiently than *TRAC*-CAR Tregs (**Fig. 3**). Further, the analysis of activation markers indicated a phenotype of *CD3ε*-TruC Tregs mirroring clinically validated polyclonal Tregs (**Suppl. Fig. 5**). Moreover, Tregs deficient of the endogenous TCR have been shown to possess reduced suppressive function^33^. Consequently, the in-depth functional characterization of gene-edited Tregs was focused on the *CD3ε*-TruC gene editing strategy.

### CD3ε-TruC Tregs upregulated ERK signaling pathway and demonstrated potent suppressive autologous and allogenic Tconv proliferation in vitro regardless of polyclonal or HLA-A2 antigen-specific stimulation

To validate signaling after TCR or antigen receptor engagement, we performed flow cytometric analysis of phosphorylated extracellular signal-regulated kinase (ERK) (**Fig. 5 A**). Prior to the functional assays *in vitro*, the expanded *CD3ε*-TruC Tregs were sorted to achieve a highly purified cell population for comparison of *CD3ε*-TruC to WT Tregs (**Suppl. Fig. 6**). Upon stimulation with plate-bound anti-CD3 mAb, *CD3ε*-TruC Tregs displayed time-dependent phosphorylation of ERK at similar extents to WT Tregs (**Fig. 5 B left; Suppl. Fig. 7**). Stimulation with plate-bound HLA-A2:Ig protein only led to ERK phosphorylation in *CD3ε*-TruC Tregs but not unedited WT Tregs (**Fig. 5 B right**).

**Figure 5.**
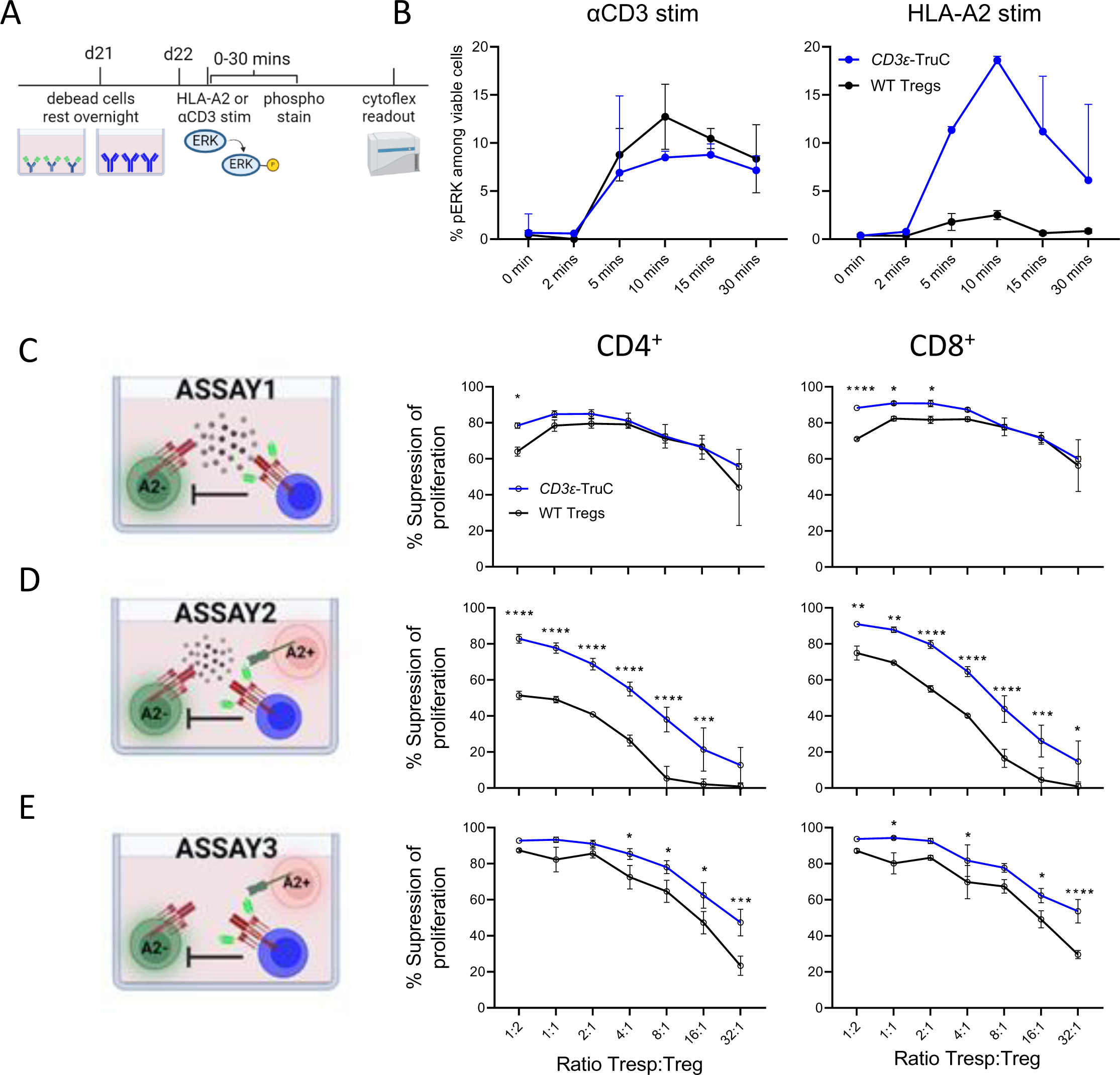
*CD3ε*-TruC Tregs upregulated ERK signaling pathway and demonstrated potent suppressive autologous and allogenic Tconv proliferation in vitro regardless of polyclonal or HLA-A2 antigen-specific stimulation. A) Experimental setup of Phosphorylation Flow assay. B) Kinetics of phosphorylation of ERK upon either 10 µg/mL αCD3 (left) or 5 µg/mL HLA-A2 (right) stimulation. N=2. Proliferation suppression % of CD4^+^ Tconv (left) and CD8^+^ Tconv (right) of *CD3ε*-TruC and WT Tregs C) upon polyclonal bead stimulation, D) upon polyclonal bead and HLA-A2*^+^* CD3-depleted PBMCs stimulation, and E) upon HLA-A2*^+^* CD3-depleted PBMCs stimulation. Data shown in C, D and E are from three technical replicates of one donor. Statistical analysis was performed using an ordinary two-way ANOVA of matched data. Multiple comparisons were performed by comparing each cell mean with the other cell mean in that row with Šidák correction. Asterisks represent adjusted *p*-values calculated in the respective statistical tests (*: *p* < 0.05; **: *p* < 0.01; ***: *p* < 0.001; ****: *p* < 0.0001).

Suppression of the unwanted proliferation of Tconv specific to auto- or allo-antigens is a major aim of Treg transfer, and this can be partially modelled *in vitro*. We used three different proliferation suppression assays for comparison of *CD3ε*-TruC Tregs and WT Tregs and evaluated the impact of the engineered antigen-specificity (**Fig. 5 C, D, E**): the first assay (1) represents the standard proliferation suppression assay using autologous responder T cells (Tresp) stimulated with anti-CD3/28 beads together with different amounts of Tregs but lacking an HLA-A2-specific stimulant. In this assay, *CD3ε*-TruC Tregs and WT Tregs suppressed the proliferation of autologous Tresp in a dose-dependent manner in all three examined donors, with more than 50% suppression capacity even at the lowest examined ratio of 1:32 (Treg:Tresp) in one donor (**Fig. 5 C, Suppl. Fig. 8 B upper plots, C upper plots**). *CD3ε*-TruC Tregs showed a significantly better Tresp proliferation suppression than WT Tregs at higher ratios of 1:2 for CD4^+^ Tresp and of 1:2, 1:1 and 2:1 for CD8 counterpart (**Fig. 5 C**). This trend was found in the second donor (**Suppl. Fig. 8 B upper plots**), interestingly not in the third donor (**Suppl. Fig. 8 C upper plots).** In the second assay (2), additional HLA-A2^+^ CD3-depleted PBMCs were added to serve as stimulant for the CAR/TruC receptors. *CD3ε-TruC* Tregs displayed a higher suppressive capacity in all three donors (**Fig. 5 D, Suppl. Fig. 8 B middle plots, C middle plots**), interestingly, WT Tregs seem to get worse when adding HLA-A2^+^ cells into the assay compared to the first assay. The third assay (3) aimed to model allogeneic T cell proliferation towards the unmatched HLA-A2^+^ CD3-depleted PBMCs and no additional anti-CD3/28 beads were included. As expected, the overall percentage of proliferating Tresp was much lower than the Tresp stimulated using anti-CD3/28 beads as stimulant (**Suppl. Fig. 8 A)**. In general, a higher proliferation suppression capacity by *CD3ε*-TruC Tregs than WT Tregs was observed in all donors except for the CD4^+^ Tresp of one donor (**Fig. 5 E, Suppl. Fig. 8 B,C lower plots**). Overall, *CD3ε-*TruC Tregs showed high suppressive capacity in autologous and allogeneic proliferation suppression assays *in vitro*.

### CD3ε-TruC Tregs efficiently prevented allogeneic cell and tissue rejection in vivo

Next, we evaluated our HLA-A2-specific *CD3ε*-TruC Tregs in a humanized *in vivo* mouse model for allogeneic cell rejection^34^. In this model, immunodeficient mice are engrafted with HLA-A2^-^ PBMCs from the same donor used for Treg engineering two weeks prior to the beginning of the experiment. Successful reconstitution was confirmed by flow cytometry (data not shown). Then, two different fluorescently labelled PBMCs were infused at a ratio of 1:1; One fraction contains autologous (auto-) PBMCs and the other fraction contains allogeneic (allo-) PBMCs from an unmatched HLA-A2^+^ donor. Co-administration of autologous Tregs should protect HLA-A2^+^ PBMCs from rejection and result into a higher allo-PBMCs to auto-PBMCs ratio at the end of the challenge (**Fig. 6 A**). The allo-PBMC to auto-PBMC ratio was measured prior to injection right after mixing as baseline. *CD3ε*-TruC Tregs significantly protected allo-PBMCs from rejection like unedited WT Tregs (**Fig. 6 B, C**). These results indicate similar suppressive capacity of *CD3ε*-TruC Tregs, comparable to the clinically-proven polyclonal Treg product^5,6^.

**Figure 6.**
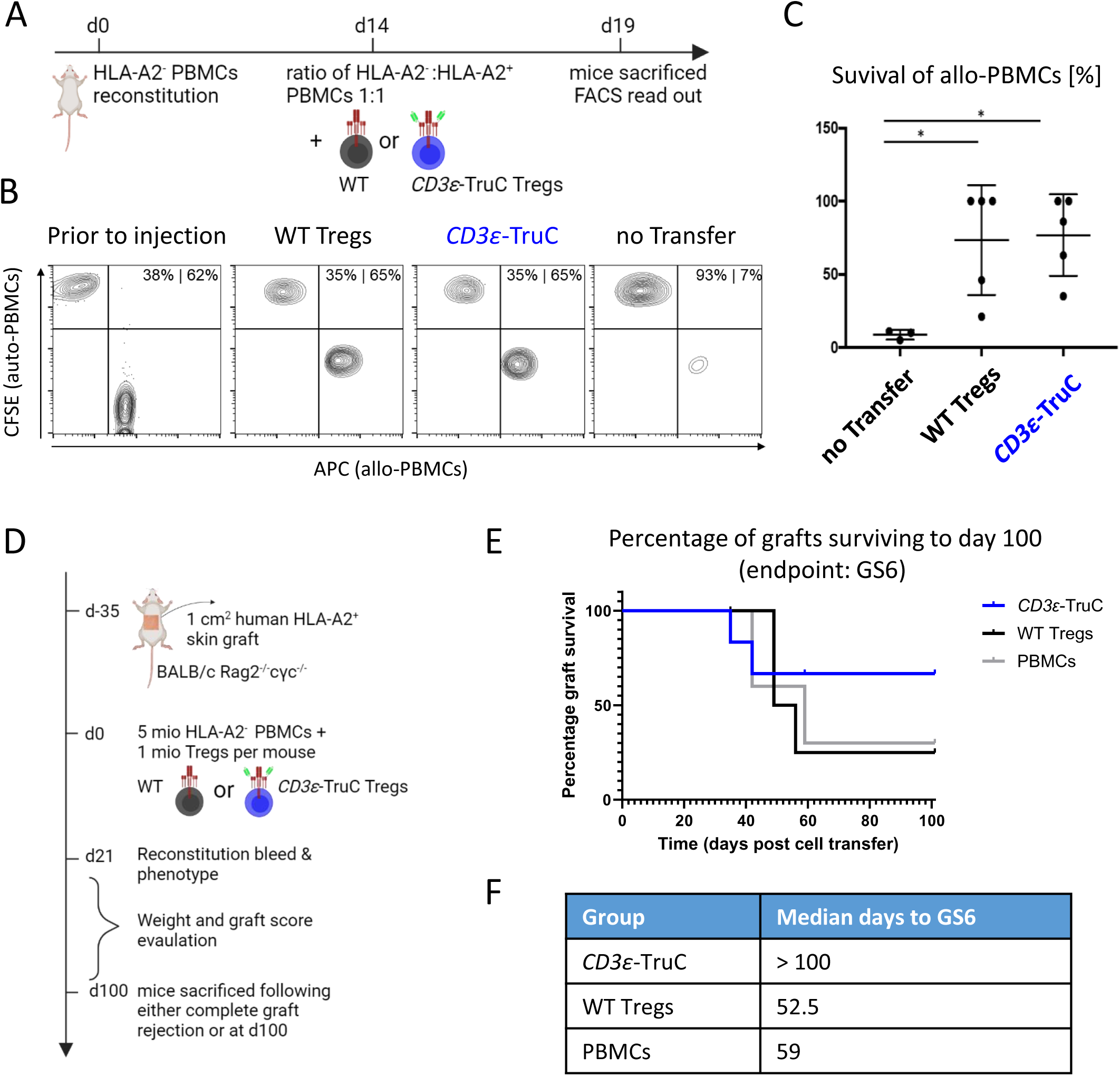
*CD3ε*-TruC Tregs efficiently prevented allogeneic cell and tissue rejection *in vivo*. A) Experimental design of a short-term allogenic donor PBMC rejection model in NSG mice *in vivo*. B) Representative FACS dot plots indicating the ratio of allogenic PBMCs to autologous PBMCs prior to injection and after injection with or without *CD3ε*-TruC Tregs or WT Tregs. C) Survival rates of allogenic PBMCs *in vivo*. N=3-5. D) Experimental setup of a long-term skin transplant model in NSG mice *in vivo*. E) Survival curves of skin graft upon allogeneic PBMCs and *CD3ε*-TruC Tregs or WT Tregs cell transfer. F) The median days to GS6 (graft scoring 6 – complete rejection) for each group are shown. N=5-6. Statistical analysis in C was performed using a one-way ANOVA of matched data. Asterisks represent adjusted *p*-values calculated in the respective statistical tests (*: *p* < 0.05).

To examine whether *CD3ε*-TruC Tregs can display improved HLA-A2-specific suppressive capacity over WT Tregs, we performed a long-term skin transplant injection model^35^ *in vivo* (**Fig. 6 D**). BALB/c Rag2^-/-^cγc^-/-^ mice were transplanted with human HLA-A2^+^ skin graft for 35 days until mice received allogenic HLA-A2^-^ PBMCs (the same donor used for Treg engineering) alone or with its autologous WT Tregs or *CD3ε*-TruC Tregs at a PBMCs:Tregs ratio of 5:1. At this high ratio of PBMCs:Tregs, polyclonal unedited WT Tregs were unable to efficiently protect the skin-allografts from rejection (**Fig. 6 E**), ending up with a median time to complete graft rejection (defined by graft score of 6, in short: GS6, see methods) of 52.5 days similar to the condition which only received the PBMCs injection. (**Fig. 6 F**). However, mice which received PBMCs and a relatively low number *CD3ε*-TruC Tregs achieved an improved long-term engraftment of the human HLA-A2^+^ skin allograft with a median time to GS6 more than 100 days (**Fig. 6 E, F**). These data collectively demonstrated the superior antigen-specific suppression capacity of *CD3ε*-TruC Tregs over unedited polyclonal WT Tregs in the long-term skin transplant model.

## IV. Discussion

This study describes an effective and GMP-compatible strategy to redirect Tregs (and other T cells) with TruC receptors using non-viral CRISPR-Cas12a gene editing of the *CD3ε* gene. Insertion of the scFv and a G4S linker into the endogenous CD3ε created TCR^+^/TruC^+^ Tregs enriched over time using anti-CD3/CD28 bead stimulation. The HLA-A2-specific gene-edited Tregs retained an epigenetic and phenotypic profile of clinically used 1^st^-generation polyclonal Treg products but benefited from the additional alloantigen-specificity during *in vitro* and *in vivo* functional tests. In the clinically relevant tissue-allograft transplantation model, a relatively low number of HLA-A2-specific *CD3ε*-TruC Tregs improved median survival of skin grafts over polyclonal Tregs. The promising data of the HLA-A2 specific *CD3ε*-TruC approach delivers a blueprint for generating redirected Treg applicable for various immunopathologies.

Non-viral gene editing is a promising approach to create genetically-modified Tregs for therapeutic applications^9^ due to economic advantages of non-viral vectors and reduced risk of insertional mutagenesis. Efficient non-viral knock-ins into Tregs has been reported previously with SpCas9^20,28,36,37^. The minimal transgene size of *CD3ε*-TruC strategy is advantageous, because linear dsDNA templates display significant dose- and also size-dependent toxicity in T cells^20,28,23,36^. Further, relative HDR rate is dependent on the targeted locus, transgene size and the indel profile of the nuclease^20,36,23,38^. AsCas12a cleaves the DNA distant of the protospacer adjacent motif and creates staggered ends; which was proposed as ideal for HDR^39^. Another potential advantage of Cas12a over Cas9 is the higher specificity^39,40^. Additionally, CAST-Seq could not detect off-target mediated translocations, suggesting high precision of the preferred crRNA-AsCas12a complex. Other unbiased assays will be needed to confirm the absence of off-target gene editing in *CD3ε*-TruC Tregs prior to clinical use.

The *CD3ε-*TruC gene editing strategy provides an elegant solution to redirect Tregs towards diverse antigens with a unified manufacturing protocol agnostic to the targeted antigen. Targeted gene insertion requires selection of a suitable locus for the respective application. Previously, *TRAC* has been the gold-standard for knock-in of 2^nd^ generation CARs^25^. Miniaturization of the CAR transgene has been proposed by inserting truncated *CD3ζ*-deficient 2^nd^-generation CARs into the *CD3ζ* gene in Tconv and Tregs^37^. Both approaches, *TRAC*^22^ and *CD3ζ*^37^, eliminate TCR surface expression in Tregs and require adaption of the manufacturing to enrich redirected Tregs via constant antigen stimuli. Similar to our approach, SpCas9-mediated *CD3ε* gene editing was used to create TruC^+^ Tconv with minimized transgenes^41,42^. In Tregs, the *CD3ε*-TruC approach enables the unique opportunity of the minimized transgene and selects for the reprogrammed Tregs by repetitive anti-CD3/CD28 stimulation applied in standard expansion protocols. In contrast, repetitive TCR/CD3-stimulation is inadvisable in Tconv due to activation-induced cell death and heightened differentiation *ex vivo*^43^, which is associated with poor effector function *in vivo*^43–46^. Safe harbor loci may also allow expression of CAR/TruC in Tregs without affecting TCR-expression, but these sites would require integration of larger transgene expression cassettes and do not enable automatic selection^47^. Essential genes, such as *GAPDH*, may be adopted for automatic selection, but fail to provide natural regulation of CAR expression^48^. Future studies may be required to evaluate whether the natural regulation of *CD3ε*-TruC provides an advantage when compared to stable overexpression.

TruC receptors exploit the endogenous TCR/CD3 complex and provide more physiological signaling to T cells^29,49^. Retrovirally-transduced TruC Tregs specific for FVIII-autoantigen demonstrated superior suppressive capacity over CAR Tregs expressing a 2^nd^-generation CAR with the CD28 costimulatory domain in a syngeneic mouse model. Using *CD3ε* gene editing, we could generate TCR^+^/TruC^+^ Tregs which responded via TCR and TruC respectively, similar to retrovirally transduced TruC Tregs^50^. *CD3ε*-TruC Tregs automatically enrich within the GMP protocols relying on anti-CD3/28 bead stimulation, enhancing the overall redirected Tregs yields in this study over TCR-deficient CAR Tregs. One potential disadvantage of TruC architecture is the lack of co-stimulatory signal by the synthetic antigen receptor. However, murine Tregs reprogrammed with 1^st^ generation CARs lacking a co-stimulatory domain displayed comparable suppressive capacity to 2^nd^ generation CAR-Tregs in syngeneic allo-transplantation models in mice^51^. Rosado-Sanchez *et al* have demonstrated that professional antigen-presenting cells can provide co-stimulation to 1^st^-generation CAR-Tregs *in trans*^51^. Therefore, *CD3ε*-TruC Tregs are likely suitable for allograft protection without the need of artificial co-stimulatory signaling in the presence of antigen-presenting cells. Future studies may investigate other co-stimulatory receptors to improve *CD3ε*-TruC Treg functionality at inflamed sites, such as the allograft.

Engineered *CD3ε*-TruC Tregs may also benefit from other modifications to improve their functionality^9^. In patients immunosuppression, Tregs could be enhanced by gene editing to induce drug resistance^52,53^ or providing artificial IL-2 cytokine signals^54,55^. Here, the minimized transgene size enables easier co-delivery of therapeutic transgenes. Further, our Cas12a editing strategy would allow combination with SpCas9-derived enzymes in a single transfection for multiplex gene editing with minimal translocations^56^. Finally, multiplex gene editing may allow to combine edits for enhanced functionality, such as drug resistance^52,53^ or *FOXP3* stabilization^57^, with modifications of HLA^58^ and enable successful off-the-shelf Treg applications.

## V. Methods and Materials

### Polyclonal Treg cell isolation and expansion

All experiments in this study were performed in accordance with the Declaration of Helsinki. Peripheral blood was obtained from healthy human adults after informed consent (Charité ethics committee approval EA4/091/19). HLA-A2 phenotype of healthy donors was determined by flow cytometry using an anti-human HLA-A2 antibody conjugated with APC (BB7.2, BioLegend) or FITC (A28, Miltenyi Biotec). Only Tregs of HLA-A2 negative donors were used for gene editing in this study. An amount of 120 mL peripheral blood was obtained from healthy donors and PBMCs were isolated using the standard density gradient centrifugation approach with Biocoll separation solution (Bio&SELL). CD4^+^ T cells were positively enriched using magnetic column enrichment with human CD4 microbeads according to the manufactureŕs recommendations (LS columns, Miltenyi Biotec). Enriched CD4^+^ T cells were stained with a cocktail of antibodies: anti-CD4 VioBlue (REAQL103), anti-CD25 APC (REAL128), anti-CD127 PE-Vio770 (REAL102), anti-CD45RA FITC (REAL164), all from Miltenyi Biotec and anti-CCR7 PE (G043H7, BioLegend). CD4^+^CD25^high^CD127^low^CCR7^+^ Tregs were sorted in a fully closed environment using a GMP-compatible Tyto sorter (Miltenyi Biotec). Sort purity of CCR7^+^ Tregs was checked on a CytoFLEX flow cytometer (Beckman Coulter) after a two consecutive sorting manner. 1×10^5^ cells were seeded in one well of a 96-well U plate (Corning) with 200 µL of Treg medium consisting of X-VIVO 15 medium (Lonza), 10% heat-inactivated fetal calf serum (FCS) (Sigma-Aldrich), 500 IU/mL recombinant human IL-2 (Miltenyi Biotec), and 100 nM rapamycin (Pfizer). All cell culture experiments were performed at a 37 °C and 5% CO2 incubator. For initial stimulation, 4×10^5^ Treg expansion beads (Miltenyi Biotec) were added to each well on the next day. Cell culture medium was replenished every 2 to 3 days. Cells were counted on day 5 and were stimulated with MACS GMP ExpAct Treg bead (Miltenyi) at a ratio of 1:1 for additional two days until electroporation.

### Conventional T cell isolation and stimulation

CD3^+^ T cells were positively enriched using magnetic column enrichment with human CD3 microbeads according to the manufactureŕs recommendations (LS columns, Miltenyi Biotec). Enriched T cells were cultured in T cell medium containing RPMI 1640 (Gibco), 10% FCS (Sigma-Aldrich), recombinant human IL-7 (10 ng/mL, Cell-Genix), and recombinant human IL-15 (5 ng/mL, Cell-Genix). Conventional T cells were stimulated via plate-bound 1 μg/mL anti-CD3 antibody (OKT3; Invitrogen) and 1 μg/mL anti-CD28 antibody (CD28.2; BioLegend) for 48 hours prior to electroporation.

### Screening of synthetic crRNAs targeted on CD3 epsilon locus by electroporation

Five guide crRNAs were selected (**Suppl. Table 1**) for screening based on the criteria of CFD off-target < 5-10 calculated by CRISPR scan^59^ and COSMID^60^. These crRNAs were synthesized from Integrated DNA Technologies (IDT). Prior to electroporation, mixing of 0.5 μL of poly(L-glutamic acid) (100 μg/μL PGA, 15-to 50-kDa, Sigma-Aldrich) with 0.48 μL of crRNA (100 uM) and 0.4 μL Alt-R A.s. Cas12a (Cpf1) Ultra (10 μg/μL = 63 μM, IDT) by thorough pipetting, with a final ratio of crRNA: Cas12a at ∼ 2:1. The mixture was incubated for 15 min at room temperature to allow for ribonucleoprotein (RNP) formation. T cells were washed twice in PBS (Gibco) and 1 ×10^6^ cells were resuspended in 20 µL of P3 buffer (Lonza). Electroporation was performed in a 16-well nucleocuvette^TM^ strip on a 4D-Nucleofector Device (Lonza) using the program EH-115. Knock-out (KO) efficiency was checked on day 4 post electroporation, and cells were stained with CD3 PacBlue (UCHT1, BioLegend) and DAPI (Thermo Fisher) and acquired on a CytoFLEX LX (Beckman Coulter). More details see Kath et al^23^.

### Generation of HDR-template for generation of TRAC-CAR and CD3ε-TruC

Four HDR-templates were used in this study, with one for integration in exon 1 of the *TRAC* locus one for integration in exon 6 of the CD3E locus, and the other two for integration in exon 3 of the *CD3Ε* locus, respectively. Anti-human HLA-A2 single chain variable fragment (scFv, clone 3PF12) was used^26^ and its cDNA sequence is available on Genbank (accession numbers AF163307 and AF163308). Anti-human HER2 scFv (clone FPR5) cDNA sequence was derived from a published patent (US8530637B2). Each donor template was designed in Snapgene (Dotmatics) and synthesized by IDT as gBlocks gene fragments with detailed nucleic acid sequences found in **Suppl. Table 2**. gBlocks were cloned in plasmid PUC19 vector backbone using multiple fragment In-Fusion cloning according to the manufactureŕs recommendation (Clontech, Takara). Plasmid transformation, colony PCR, and plasmid purification and validation were previously described^23^. HA-flanked HDR-templates were amplified from the validated plasmids by PCR using the KAPA HiFi HotStart 2x Readymix (Roche). Primers used in this study can be found in **Suppl. Table 3**. PCR products were purified and concentrated using paramagnetic beads (AMPure XP, Beckman Coulter Genomics). The concentration of HDR-templates was determined by using a NanoDrop 1000 spectrophotometer (ThermoFisher) and a Qubit Fluorometer (ThermoFisher). HDR-templates were adjusted to a concentration of 1 μg/μL in nuclease-free water and stored at -20°C until use.

### Gene editing of Tregs with HDR-templates via electroporation

Upon 7 to 9 days of stimulation of Tregs, beads were removed by a MACSiMAG^TM^ separator (Miltenyi Biotec) and bead-free Tregs were washed in PBS and suspended in P3 buffer as conventional T cells. 0.5 µg of *TRAC*-CAR HDR-template or *CD3ε*-TruC HDR-template was used to mix with RNP and cell suspension prior to electroporation. Same as conventional T cells, electroporation was performed on a 4D-Nucleofector Device (Lonza) using the program EH-115. To rescue cells, 90 µL of pre-warmed Treg medium was added to each well immediately after electroporation, incubated for 10 mins in a 37°C incubator prior to evenly transferring them to 2 wells of a 96-well U plate containing 150 µL pre-warmed Treg medium. On the next morning, cells were stimulated with Treg expansion beads at a ratio of 1:1.

### CAST-Seq analysis

Genomic DNA was extracted using the NucleoSpin Tissue® kit (Macherey-Nagel). CAST-Seq analyses were performed following the previously described protocol^32^ with some adjustments to the workflow^61^. In brief, the average fragmentation size of the genomic DNA was aimed at a length of 500 bp. The libraries were sequenced on a NovaSeq 6000 using 2 × 150 bp paired-end sequencing (GENEWIZ, Azenta Life Sciences). Changes were made to the bioinformatic pipeline to enhance specificity. Sites under investigation were categorized as OMT if the *p*-value met the cutoff of 0.005. In addition, further features were incorporated in the CAST-Seq algorithm, including barcode hopping annotation and an update to the coverage analysis to reduce the execution time by aligning the gRNAs only to the most covered regions for each site. Two technical replicates from samples of two different donors were used for the CAST-Seq analysis. Only sites identified as significant hits in both biological replicates were considered as putative events. Results are showed in Figure 2A and **Suppl. Table 4**.

### T7E1 Assay

Briefly, 50 ng of genomic DNA was subjected to PCR reaction and 100 ng of the resulting amplicons were subjected to a slow reannealing for 5 minutes at 95°. The product was then digested with T7E1 (NEB) at 37°C for 20 minutes. and then resolved through a 2% agarose gel electrophoresis.

### Expansion of CD3ε-TruC and TRAC-CAR Tregs

*TRAC*-CAR Tregs were initially stimulated with either 1) plate-bound 2 µg/mL HLA-A2:Ig fusion protein (BD), or 2) plate-bound 2 µg/mL HLA-A2:Ig fusion protein (BD) with plate-bound 1 µg/mL anti-CD28 (BioLegend), or 3) gamma-irradiated (30 Gy) HLA-A2^+^ K562 cell line (a gift from Fatih Noyan, MHH, Germany) at a ratio of 1:1. Stimulation and medium replenishment were repeated every 2 to 3 days. Subsequently, *CD3ε*-TruC and *TRAC*-CAR Tregs were expanded with Treg expansion beads (Miltenyi Biotec) every 2 to 3 days at a cell-bead ratio of 1:1 for two weeks post electroporation. Beads were removed and bead-free Tregs rested overnight prior to *in vitro* assay setup.

### Gene editing efficiency check by flow cytometry

*TRAC*-CAR Tregs were stained with anti-CD3 PB (UCHT1, BioLegend) and anti-Fc A647 (Jackson Immuno Research Labs). *CD3ε*-TruC Tregs were stained with CD3 PB and anti-Myc A647 (9B11, Cell Signaling). Cells were resuspended in DAPI containing PBS prior to acquisition by CytoFLEX LX. Gene editing efficiency check of Tregs were performed at day 11, day 15 and day 22 upon blood withdrawal.

### Phenotypic characterization of gene-edited Tregs

Tregs were stained with fixable dye Aqua (Invitrogen), were then fixed and permeabilized using Transcription Factor Buffer Set (BD). Anti-Fc A647 for *TRAC*-CAR Tregs and anti-Myc A647 for *CD3ε*-TruC Tregs were stained intracellularly, followed by 2 times wash using 1× permeabilization buffer (BioLegend). Cells were then stained by an antibody cocktail of anti-CD3 PB, BioLegend, anti-CD4 PE (13B8.2, Beckman Coulter), anti-CD25 PC7 (B1.49.9, Beckman Coulter), and anti-FOXP3 FITC (259D/C7, BD). For cytokine profile examination of gene-edited Tregs, 3x10^5^ cells were stimulated by 10 ng/mL PMA and 2.5 µg/mL Ionomycin (Sigma Aldrich) for 6 hours. Brefeldin A (Sigma Aldrich) was added at a concentration of 10 µg/mL after 1 h of co-culture. Upon 6 hour of stimulation, cells were harvested and stained intracellularly with anti-TNFα A700 (Mab11, BioLegend), anti-IFNγ APC-eF780 (4S.B3, Invitrogen), and anti-IL-2 PE-Cy7 (MQ1-17H12, BioLegend). Stained cells were acquired at Cytoflex LX.

### DNA methylation analysis of the FOXP3 TSDR using bisulfite amplicon sequencing

Amplicon TSDR methylation analysis was performed as previously described^62^. Briefly, 1x10^6^ cells were snap-frozen and stored at liquid nitrogen until genomic DNA isolation with the Quick-DNA Microprep Kit (Zymo Research D3020) according to the manufactureŕs protocol. 200 ng of genomic DNA was bisulfite-converted using an EZ-DNA methylation Gold kit (Zymo Research D5006).

PCRs were performed with input of 200 ng of bisulfite-treated DNA, 2 x KAPA HiFi Hotstart Uracil+ ReadyMix (Kapa Biosystems KK2802), 0.25 mM of each dNTP, 0.3 pmol of primers (F1: 5’ ACACTCTTTCCCTACACGACGCTCTTCCGATCTTTTGGGGGTAGAGGATTTAGA GGG-3’ and R3: 5’GACTGGAGTTCAGACGTGTGCTCTTCCGATCTCCACATCCACCAACACCCAT -3’) with Illumina compatible universal adaptor sequences attached at the 5’-end. Amplicons were purified with DNA Clean & Concentrator-5 Kit (Zymo Research D5006) and normalized to 20 ng/uL for sequencing by Genewiz/ Azenta aiming at 1x10^4^ reads per amplicon. Reads were aligned and evaluated using the Bismark package^63^.

### Activation profiles of CD3ε-TruC versus TRAC-CAR Tregs

Tregs were rested overnight without IL-2, rapamycin and beads. Plate coating of HLA-A2: Ig dimer (BD) in a serial dilution of 10 - 5 - 2.5 - 1 μg/mL with 1 μg/mL anti-CD28 antibody in a 96-well flat plate was prepared one day in advance. 4x10^5^ *CD3ε*-TruC and *TRAC*-CAR as well as WT Tregs were stimulated for 24 hours, respectively. Upon 24 hours stimulation, cells were stained with antibodies: anti-CD25 PC7 (B1.49.9, Beckman Coulter), anti-CD71 BV786 (M-A712, BD bioscience), anti-FOXP3 FITC (259D/C7, BD), anti-ICOS BV650 (C398.4A), anti-LAP PE (S20006A), anti-CTLA-4 BV421 (BNI3), anti-CD137 PE (4B4-1), anti-CD154 BV421 (24-31), anti-Helios Percp-Cy5.5 (22F6) (all from BioLegend). Among them, anti-Helios and anti-FOXP3 antibodies were stained intracellularly.

### Purification of gene-edited Tregs using a Tyto sorter

*CD3ε*-TruC Tregs were sorted as Myc^+^ using a Tyto sorter. Sorting was performed two rounds to achieve a high purity. Sorted cells were further expanded with Treg expansion beads and were used for *in vitro* Phosphorylation assay and suppression assay.

### Phosphorylation of ERK upon stimulation

Plate coating of either 5 μg/mL HLA-A2:Ig dimer (BD) or 10 μg/mL anti-CD3 antibody (Invitrogen) was prepared one day in advance. Tregs were firstly stained with fixable live/dead dye UV (Invitrogen). 5x10^5^ cells were seeded to each well by a short spin down, then immediately incubated at 37°C on a thermomixer (Eppendorf). Upon indicated time period: 0 min, 2 mins, 5 mins, 10 mins, 15 mins, and 30 mins, 200 μL of pre-warmed 4% fixation buffer (BioLegend) was added to cells for additional 15 minutes incubation. Cells were then washed with 200 μL of True-Phos perm buffer (pre-cooled at -20°C, BioLegend) and incubated at -20°C for 60 min. Cells were washed twice with PBS prior to intercellular stain with anti-ERK1/2 Phospho (Thr202/Tyr204) FITC (6B8B69, BioLegend) for 30 minutes at 4°C. Cells were washed twice with PBS and acquired at Cytoflex LX.

### Suppression assay

C) upon polyclonal bead stimulation, D) upon polyclonal bead and HLA-A2*^+^* CD3-depleted PBMCs stimulation, and E) upon HLA-A2*^+^* CD3-depleted PBMCs stimulation. Autologous Tconv cells were enriched from PBMCs as responder cells (Tresp). Tresp were labeled with 1 µM of CFSE (Thermo Fisher) for 10 minutes at 37°C. 5x10^4^ labeled Tconv were seeded in each well of a 96-well U plate. Tregs were seeded at various ratios of Tresp to Treg ranging from 1:2 to 32:1. Three stimuli conditions were tested in this study: 1) using Treg suppression inspector, human (beads, Miltenyi Biotec) at a 1:1 ratio of total seeded cells per well to beads; 2) using polyclonal beads and HLA-A2^+^ CD3-depleted PBMCs that werepre-labeled with CT FarRed (Thermo Fisher) at a 1:1:1 ratio to Tresp + Tregs; 3) using HLA-A2+ CD3-depleted PBMCs at a ratio of 1:1 to Tresp + Tregs. Each condition has technical triplicates. Wells containing Tresp with stimulus but not Tregs were served as positive control of proliferation. Wells containing Tresp alone were served as negative control of proliferation. Cells were all resuspended in X-VIVO 15 medium supplemented with 10% FCS and 1% Penicillin/Streptomycin. Upon 5 days of co-culture, cells were harvested and stained with anti-CD4 PE (13B8.2, Beckman Coulter) and anti-CD8 PE-Cy7 (RPA-T8, BD Pharmingen) and resuspended in DAPI containing PBS for immediate acquisition at Cytoflex LX. % suppression capacity of Tregs = (% divided Tresp alone - % divided Tresp treated with Tregs)/ % divided Tresp alone *100.

### Allogeneic rejection model

The *in vivo* functionality of genetically edited Tregs was evaluated using an allogeneic rejection model in NOD.Cg-Rag1^tm1Mom^Il2rγ^tm1Wjl^/SzJZtm (NRG) mice, as described by Henschel et al.^34^. NRG mice were bred in a pathogen-free environment at the animal facility of Hannover Medical School (Hannover, Germany). All animal procedures were approved by the local Animal Ethics Review Board (Oldenburg, Germany, 21/3621) and conducted in compliance with prevailing German regulations. NRG mice were reconstituted with 5x10^6^ autologous HLA-A2*02:01-negative PBMCs. Successful reconstitution was confirmed via whole blood analysis obtained from the submandibular vein after two weeks, using staining with anti-hCD4 BV421 (RPA-T4; BioLegend) and anti-hCD8 APC (SK1; BioLegend). On Day 14, mice subjected to the in vivo killing experiments were intravenously injected with 5×10^5^ allogeneic HLA-A02:01-positive PBMCs labeled with eF670 (5 µM, CellTrace, ThermoFisher) and 5×10^5^ syngeneic HLA-A02:01-negative PBMCs labeled with CFSE (0.5 µM, CellTrace, ThermoFisher) at a 1:1 ratio, along with 5 ×10^5^ WT or *CD3ε*-TruC Tregs. Five days post-transfer, splenic cells were isolated, and the survival of allogeneic target cells was assessed via flow cytometry.

### Humanized skin allograft mouse model

The skin allograft model was performed as previously described^35^. Briefly, BALB/cRag2^-/-^cγc^-/-^ mice (Jackson Laboratory, Bar Harbor, ME, USA) were housed in individually ventilated cages under specific pathogen-free conditions. All protocols were conducted in accordance with the UK Animals (Scientific Procedures) Act (1986) and approved by Oxford University’s Committee on Animal Care and Ethical Review. Human skin was procured with full informed written consent and with ethical approval from the Oxfordshire Research Ethics Committee (REC B), study number 07/H0605/130. 1 cm^2^ HLA-A2 positive human skin graft was surgically grafted onto the flank of BALB/cRag2^-/-^cγc^-/-^ recipient. Age- and sex-matched mice were used. Five weeks following skin grafting, mice received 5x10^6^ cryopreserved PBMCs (same as Treg donor) in pure RPMI 1640 medium via intra-peritoneal injection, with or without 1x10^6^ *in vitro*-expanded WT Tregs or HLA-A2-specific *CD3ε*-TruC Tregs. Grafts were observed for macroscopic markers of rejection by an assessor blinded to treatment groups. Mice were sacrificed following either graft rejection or at day 100 post-transplantation by cervical dislocation.

### Data analysis, statistics

Flow cytometry data were analyzed with FlowJo software v.10 (BD). Graphs and statistical analyses were created using Prism 9 (GraphPad). T7E1 indel % was calculated by ImageJ analysis. Schemes and experimental setup layout were created with BioRender.com.

## VI Acknowledgements

We would like to express our gratitude to the following individuals for their valuable contributions: Anne Schulze from Julia Polansky group (BIH, Berlin, Germany) for her technical assistance with the TSDR experiment presented in Fig. 4 E. Sarah Schulenberg from Michael Schmueck-Henneresse group (Charité, Berlin, Germany) for sharing her phosphorylation flow cytometry protocol. Cell sorting was performed using the Tyto sorter instrument at the BIH Core Unit for pluripotent Stem Cells and Organoids (BIH-CUSCO). Geoffroy Andrieux (from the Institute of Medical Bioinformatics and Systems Medicine, Medical Center-University of Freiburg) for his help with the bioinformatic part in the CAST-Seq pipeline. This project has received funding from the European Union’s Horizon research and innovation program under grant agreements no. 825392 (ReSHAPE) and no. 101057438 (geneTIGA) and by the European Research Council under the ERC starting grant EpiTune (Grant Agreement Nr. 803992 to J.K.P.). Views and opinions expressed are however those of the author(s) only and do not necessarily reflect those of the European Union, the European Health and Digital Executive Agency (HADEA) or the ERC. Neither the European Union nor the granting authority can be held responsible for them.

## VII. Author contributions

W.D. designed parts of the study, planned, and performed experiments, analyzed results, interpreted the data, and wrote the manuscript. F.N., O.M., V.D., V.G., C.F.-G., planned and performed experiments, analyzed results, interpreted the data, and edited the manuscript. J.K. designed HLA-A2-specific CARs and HER2 CAR construct, performed experiments, interpreted data and edited the manuscript. M.Y., M.S. performed experiments and analyzed results and edited the manuscript. O.W. provided reagents, interpreted results and edited the manuscript. J.K.P., T.C., J.H., F.I., E.J., H.-D.V., P.R., M.S.-H. supervised parts of the study, provided reagents, interpreted data and edited the manuscript. D.L.W. designed and led the study, planned experiments, analyzed results, interpreted data, and wrote the manuscript. All authors reviewed and approved the manuscript in its final form.

## VIII. Conflict of Interest Disclosures

W.D, J.K., H.-D.V., P.R., M.S.-H. and D.L.W. are listed as inventors on a patent application related to the work presented in this manuscript. H.-D.V. is co-founder and CSO at CheckImmune GmbH. P.R., H.-D.V., O.W. and D.L.W. are co-founders of the startup TCBalance Biopharmaceuticals GmbH focused on regulatory T cell therapy. All other co-authors report no conflict of interest related to this work.

