## Supplemental Figures for "Gene editing of *CD3 epsilon* gene to redirect regulatory T cells for adoptive T cell transfer"

**Summary**

**Supplemental Figures**

**Suppl. Fig.1 *TRAC* replaced HLA-A2 specific CAR Tregs were successfully generated via non-viral gene editing but suffered from poor expansion.**

**Suppl. Fig. 2 Highly purified polyclonal Tregs were achieved after two consecutive sorting.**

**Suppl. Fig. 3 T7E1 assay and Sanger sequencing confirmed high cutting efficiency of Cas12a-crRNA #1 targeting *CD3ε* exon 3.**

**Suppl. Fig. 4 Comparison of HLA-A2 scFv knock-in efficiency in Tconv and Tregs.**

**Suppl. Fig. 5 Activation profiles of *CD3ε*-TruC Tregs compared to *TRAC*-CAR and WT Tregs upon antigen HLA-A2 specific stimulation.**

**Suppl. Fig. 6 Highly purified *CD3ε*-TruC$^+$ Tregs were achieved for functional assays *in vitro*.**

**Suppl. Fig. 7 HLA-A2-specific *CD3ε*-TruC Tregs showed time-dependent phosphorylation of ERK upon antigen stimulation.**

**Suppl. Fig. 8 Histogram plots of autologous Tconv proliferation without Treg co-culture.**

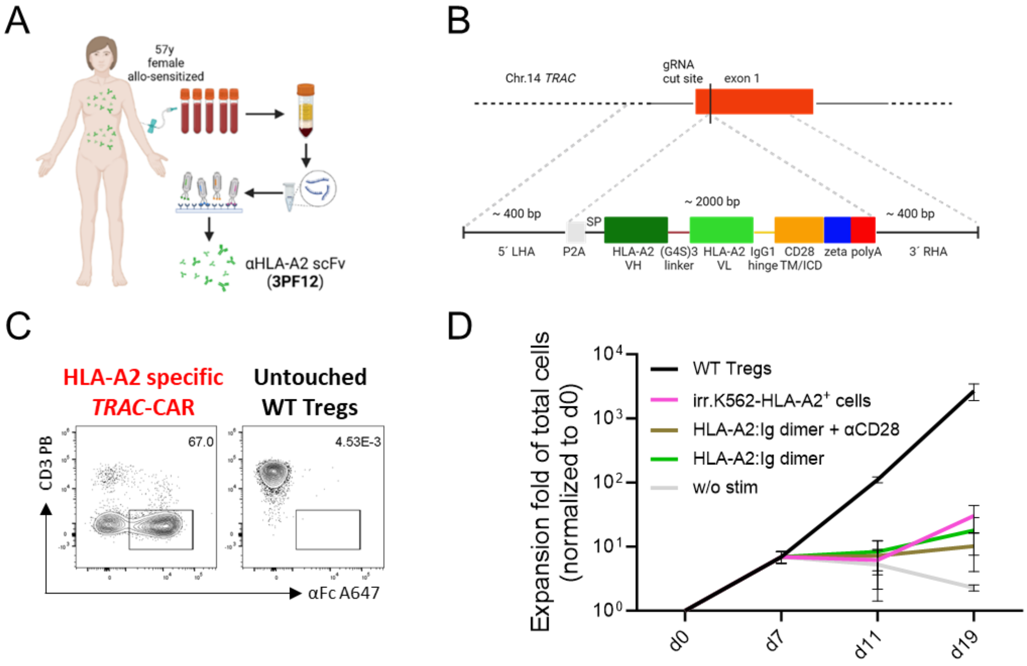


**Suppl. Fig. 1 *TRAC* replaced HLA-A2 specific CAR Tregs were successfully generated via non-viral gene editing but suffered from poor expansion.** A) Anti-human HLA-A2 scFv clone 3PF12 with medium affinity[26] was used in this study. B) HDR-mediated integration of a HDR-template into *TRAC* locus. HDR-template contains a 400 bp left homology arm (LHA), self-cleaving peptide P2A, signal peptide (SP), heavy chain and light chain of HLA-A2 scFv with 3 × G4S linker in between, IgG1 hinge (used for detecting CAR integration), CD28 transmembrane domain (TD) and intracellular domain (ICD), CD3 zeta chain, followed by a poly (A) tail and 400 bp right homology arm (RHA). C) Representative dot plots of αFc A647 against αCD3 PB stained on HLA-A2 specific *TRAC*-CAR Tregs and untouched wild type (WT) Tregs. Cells were pre-gated on live single lymphocytes. D) Expansion fold change during 2-week expansion post electroporation of *TRAC*-CAR Tregs upon either irradiated K562.HLA-A2$^{+}$ cells (ratio 1:1), or plate-coated 5 µg/mL HLA-A2:Ig dimer with or without 1 µg/mL αCD28, or without stimulation. WT Tregs with Treg expansion bead stimulation (ratio 1:1) were used as positive control. Tregs were stimulated every 2 to 3 days. N=3.

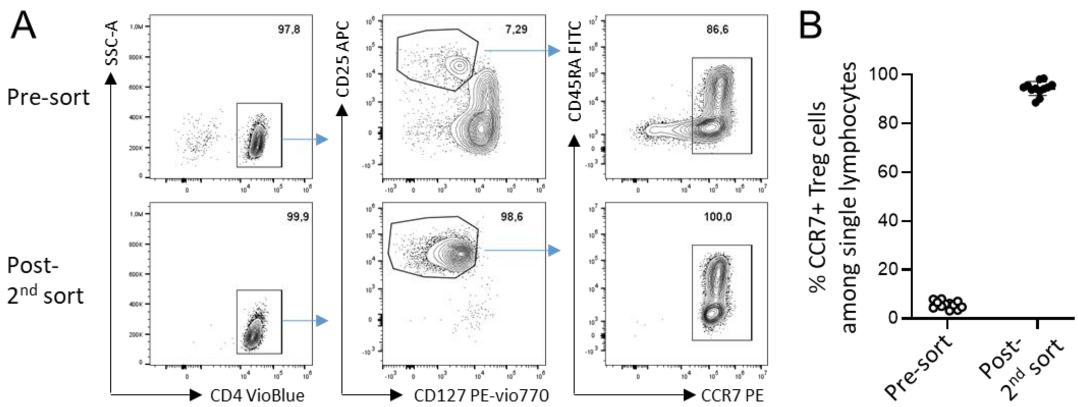


**Suppl. Fig. 2 Highly purified polyclonal Tregs were achieved after two consecutive sorts.** A) Representative dot plots of $CD4^{+}CD25^{high}CD127^{low}CCR7^{+}$ Tregs pre-sort (upper) and post two consecutive sort by a Tyto sorter (lower). Cells were pre-gated on single viable lymphocytes. B) Percentages of $CCR7^{+}$ Tregs among single lymphocytes pre- and post-$2^{nd}$ sort. N=7.

A

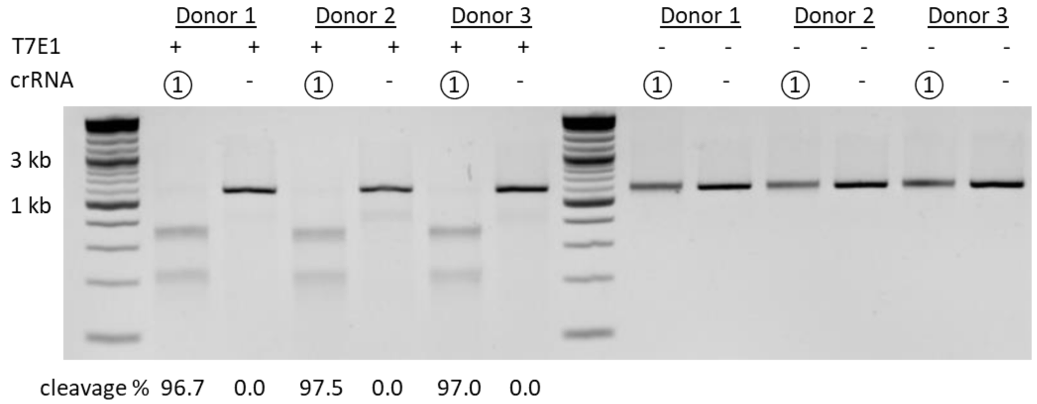


B

| Samples | | T7E1 cleavage % | Indel % $\%\ \boldsymbol{edited} = \mathbf{100} \times (\mathbf{1} - \sqrt{(\mathbf{1} - \boldsymbol{fraction\ cleaved})})$ |
|---|---|---|---|
| Donor 1 | KO | 96.7 | 81.8 |
| | mock | 0.0 | — |
| Donor 2 | KO | 97.5 | 84.2 |
| | mock | 0.0 | — |
| Donor 3 | KO | 97.0 | 82.7 |
| | mock | 0.0 | — |

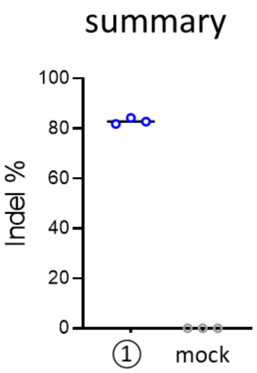


C

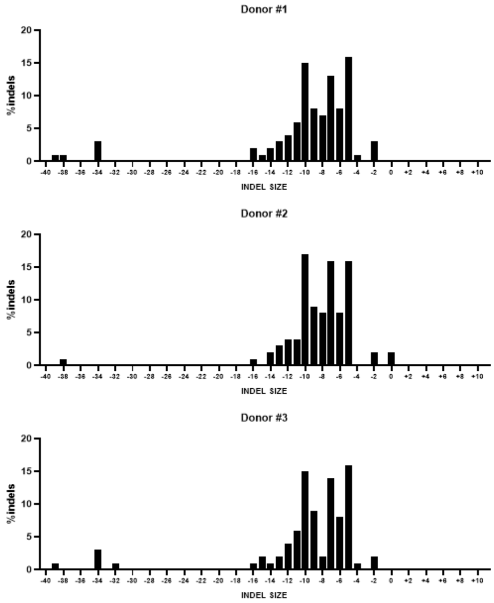


**Suppl. Fig. 3 T7E1 assay and Sanger sequencing confirmed high cutting efficiency of Cas12a-crRNA #1 targeting *CD3ε* exon 3.** A) Electrophoresis gel image and B) T7E1 cleavage % and Indel % summary of genomic DNA of crRNA #1 targeted *CD3ε*-KO Tconv and mock Tconv upon T7 Endonuclease I treatment. C) ICE analysis of *CD3ε*-KO Tconv indicating indel contributions. N=3.

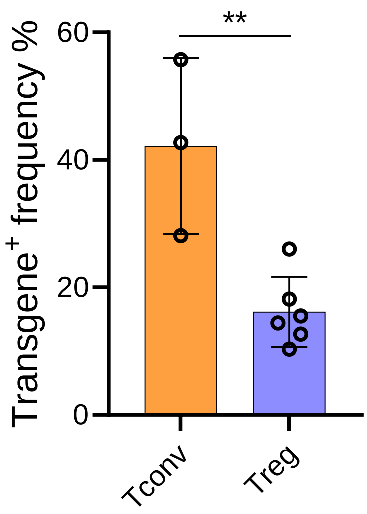


**Suppl. Fig. 4 Comparison of HLA-A2 scFv knock-in efficiency in Tconv and Tregs.** N=3 for Tconv (yellow). N=6 for Tregs (light blue). Statistical analysis was performed using a t-tests of unpaired data. Asterisks represent two-tailed *p*-value (**: $p < 0.01$).

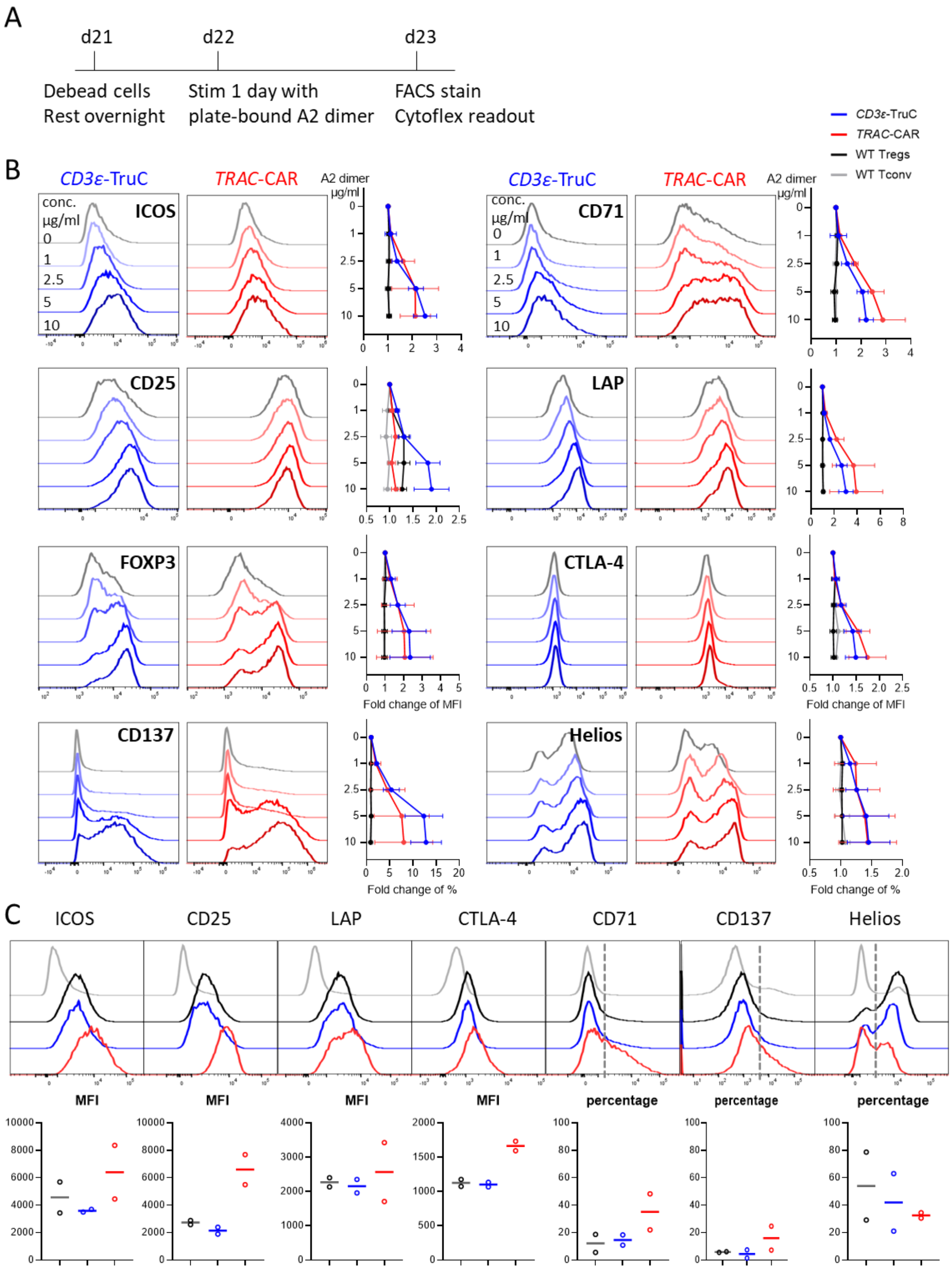


**Suppl. Fig. 5 Activation profiles of *CD3ε*-TruC Tregs compared to *TRAC*-CAR and WT Tregs upon antigen HLA-A2 specific stimulation.** A) Experimental setup of activation profile assay. B) Representative offset histograms and summary of each investigated activation marker of *CD3ε*-TruC (blue) and *TRAC*-CAR (red) upon a gradient concentration of plate coated HLA-A2:Ig fusion protein stimulation. C) Representative offset histograms (upper) and summary (lower) of activation markers of *CD3ε*-TruC (blue) and *TRAC*-CAR (red) compared to WT Tregs (dark) without stimulation for 1 day. WT Tconv (light gray) used as negative control. N=2.

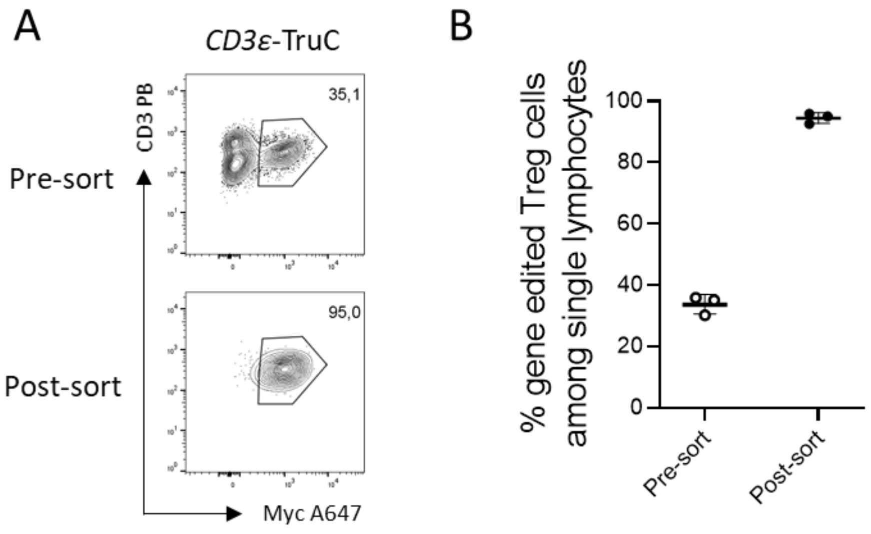


**Suppl. Figure 6 Highly purified *CD3ε*-TruC⁺ Tregs were achieved for functional assays *in vitro*.** A) Representative dot plots of CD3⁺Myc+ *CD3ε*-TruC Tregs pre-sort (upper) and post-sort (lower) using a Tyto sorter. Cells were pre-gated on single viable lymphocytes. B) Percentages of *CD3ε*-TruC⁺ Tregs among total viable Tregs pre- and post-sort from three biological donors. N=3.

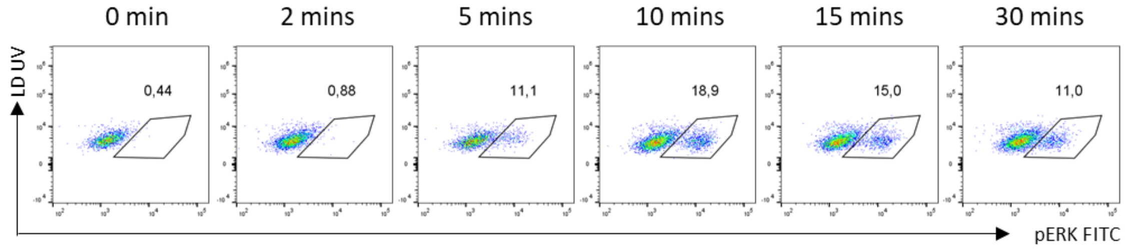


**Suppl. Figure 7 HLA-A2-specific *CD3ε*-TruC Tregs showed time-dependent phosphorylation of ERK upon antigen stimulation.** Representative FACS plots showing proportion of phosphorylated ERK among viable cells upon plate-bound 5 µg/mL HLA-A2:Ig fusion protein stimulation.

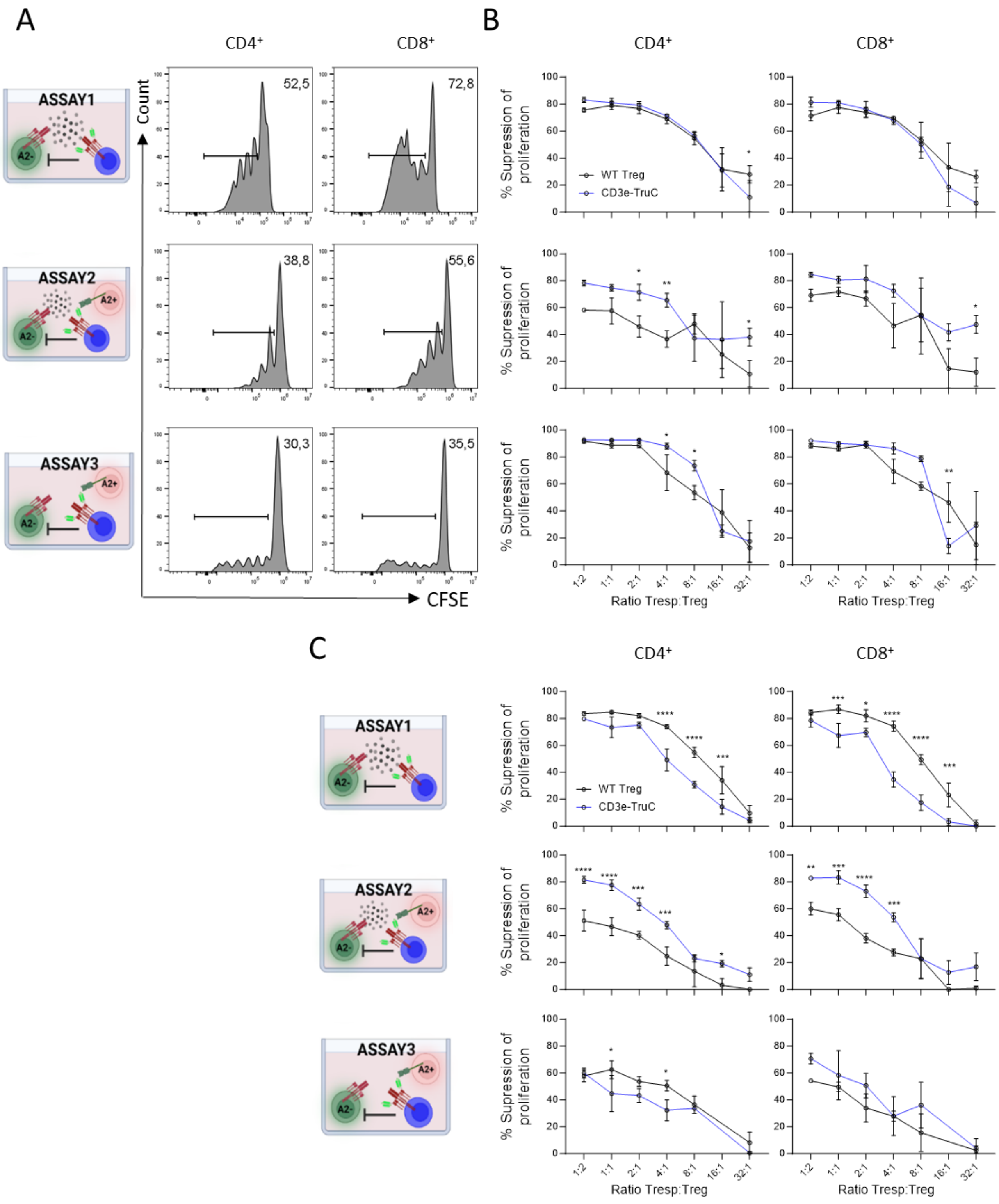


**Suppl. Fig. 8 *CD3ε*-TruC Tregs suppressed autologous and allogenic Tconv proliferation functionally *in vitro*.** A) Histogram plots of autologous and allogenic Tconv proliferation without Tregs co-culture. CD4$^+$ (left) and CD8$^+$ (right) Tconv cells were co-culture with either B, C top) MACS GMP ExpAct Treg bead at a ratio of 1:1 or B, C middle) MACS GMP ExpAct Treg bead together with HLA-A2$^+$ CD3 depleted PBMCs at a ratio of 1:1:1 or B, C bottom) HLA-A2$^+$ CD3 depleted PBMCs at a ratio of 1:1. Data shown in B and C are from three technical replicates of two donors, respectively.  Statistical analysis was performed using an ordinary two-way ANOVA of matched data. Multiple comparisons were performed by comparing each cell mean with the other cell mean in that row with Šidák correction. Asterisks represent adjusted *p*-values calculated in the respective statistical tests (*: $p < 0.05$; **: $p < 0.01$; ***: $p < 0.001$; ****: $p < 0.0001$).
